# Free acid β-hydroxybutyrate supplementation does not ameliorate dextran sodium sulfate-induced colitis similar to ketogenic diet in male mice

**DOI:** 10.1101/2025.01.19.633758

**Authors:** Lotta Toivio, Jyri Toivio, Jere Lindén, Keehoon Lee, Markku Lehto, Hanne Salmenkari, Riitta Korpela

## Abstract

High-fat, low-carbohydrate ketogenic diets have been found to alleviate experimental colitis in rodents. These diets lead to increased endogenous production and utilization of ketone bodies, such as β-hydroxybutyrate (BHB), and supplementation with exogenous ketones has arisen as a potential alternative to ketogenic diets. This study aimed to investigate how continuous high-dose feeding with free acid BHB influences experimental colitis compared to a ketogenic diet with the hypothesis that BHB would also alleviate the inflammation. We fed nine-week-old C57BL/6J male mice for four weeks with one of three diets: a low-fat control diet, a ketogenic diet, or a low-fat diet supplemented with free acid R-BHB and then induced colonic inflammation with dextran sodium sulfate (DSS). We assessed macroscopic and histological changes in the colon, intestinal permeability to fluorescein isothiocyanate dextran, colonic mRNA expression of tight junction proteins and inflammatory markers, fecal calprotectin, and microbiota composition. While the ketogenic diet alleviated DSS-induced weight loss, macroscopic changes, and histological lesions, the BHB-supplemented diet did not have the same effect. The pre-DSS composition of the microbiota was drastically different between the diet groups which may partly explain the different outcomes. In conclusion, high-dose supplementation with free acid BHB may not produce the same benefits as ketogenic diet in the context of colonic inflammation.

## 1. Introduction

Inflammatory bowel diseases (IBDs) are chronic, debilitating conditions of the intestine to which there is no known cure. IBDs comprise of Crohn’s disease and ulcerative colitis, and the clinical presentations of these diseases include abdominal pain, diarrhea, rectal bleeding, and weight loss. In the current care, IBDs are treated with pharmacotherapies, such as aminosalicylates, corticosteroids, immunomodulators, and biologics, especially tumor necrosis alpha-inhibitors (Cai et al., 2021). However, a substantial number of patients fails to respond to these therapies (Alsoud et al., 2021) which calls for novel treatment strategies.

High-fat, low-carbohydrate ketogenic diets have arisen as potential adjunct therapies for chronic diseases (Choi et al., 2020; Watanabe et al., 2020). On such a diet, ketogenesis is upregulated in the liver due to the decreased availability of glucose, which leads to increased circulating levels of ketone bodies acetoacetate and β-hydroxybutyrate (BHB). Most tissues of the body, including the intestine (Windmueller & Spaeth, 1978), can use these metabolites as an energy source. In addition to providing energy, BHB has anti-inflammatory properties (Shippy et al., 2020) and it can decrease oxidative stress (Shimazu et al., 2013). Recently, supplementation with exogenous ketone bodies has emerged as an alternative strategy to increase ketone levels without carbohydrate restriction (Yu et al., 2023).

Previously, we and others have observed ketogenic diets to ameliorate experimental colitis in rodents and it seems that the effect of the diet is mediated through alterations in the intestinal microbiota (Abdelhady et al., 2023; Kong et al., 2021; Toivio et al., 2024). One mechanism through which these diets may modify the microbiota is via ketone bodies, specifically BHB which is also metabolized by the gut microbiota (Sasaki et al., 2020). BHB appears to alter gut microbiota in a way that reduces pro-inflammatory Th17 cells in the intestine. In line with this, previous studies have shown BHB to alleviate dextran sodium sulfate (DSS)-induced colitis in mice when administered rectally (Huang et al., 2022), via a gastric gavage (Li et al., 2021), or intraperitoneally (Abdelhady et al., 2023; Mohammed et al., 2024). Injected BHB was reported to abate DSS-induced changes in the microbiota in a similar fashion as a ketogenic diet (Abdelhady et al., 2023). In addition, continuous feeding with a ketone monoester exhibited a pronounced benefit over intraperitoneal BHB injection on colitis in rats (Mohammed et al., 2024), indicating that the effects of ketone bodies in the gut lumen, such as interaction with the gut microbiota, may be of importance. As a treatment for chronic diseases such as IBDs, ketone body supplementation would be more feasible and sustainable when compared to a ketogenic diet and thus, warrants further research. However, besides the increased supply of ketone bodies, other features of ketogenic diets such as the macronutrient composition, can also be responsible for the diet’s effect on microbiota and thus, the alleviation of colitis.

As previous studies have not compared the effects of oral BHB supplementation and ketogenic diets on colitis, we set out to investigate whether BHB administration would alleviate colonic inflammation alike the diet. To investigate this, we fed mice a low-fat diet either with or without free acid R-BHB, the bioidentical form of the compound, or a ketogenic diet for four weeks, after which colitis was induced with DSS. We compared macroscopic and histological changes in the colon, fecal calprotectin, colonic mRNA expression of inflammatory markers and tight junction (TJ) proteins, intestinal permeability, and microbiota composition with the aim to clarify if continuous high-dose feeding with BHB recapitulates the beneficial effects of ketogenic diets in experimental colitis.

## 2. Materials and methods

### 2.1. Animal experiment

The experiment was approved by the animal research board of the Regional State Administrative Agency for Southern Finland (ESAVI/9377/2019) and conducted according tho the ARRIVE guidelines (Percie du Sert et al., 2020). Eight-week-old male C57BL/6J mice (n = 44) were obtained from Scanbur (Karlslunde, Denmark) and allowed to acclimatize for seven days before the start of the dietary interventions. The animals were housed in individual cages under a 12 h light–dark cycle, at 25 ± 1 °C and 50–60 % humidity with unrestricted access to food and water. Sample size was determined based on our previous experience on the DSS model of experimental colitis (Toivio et al., 2024).

The mice were randomly divided into three groups based on the diet: low-fat control diet group (CD) (n = 16), ketogenic diet group (KD) (n = 14), and BHB-supplemented diet group (BHB) (n = 14). One animal in KD had to be euthanized after three weeks of the intervention due to progressive weight loss. Weight and consumption of food and fluid were measured daily. The study diets were custom-made (Envigo, Indianapolis, IN, USA) and matched for protein and micronutrients. Free R-BHB incorporated into the diet was supplied by NNB Nutrition (Nanjing, China). The diet formulations are provided in Table 1 and macronutrient compositions of the diets in Table 2. The percentage of BHB (12 E%), accounting in the diet was determined based on previous literature (Poff et al., 2014; Yurista et al., 2021). In human studies, doses up to 752 mg/kg of BHB, possibly accounting for 8-12 % of daily energy intake, as ketone esters have been tested without serious adverse effects (Dearlove et al., 2021). Duration of the experiment was based on our previous research using ketogenic diets in the same model (Toivio et al., 2024).

**Table 1.** Formulations of study diets.

| Ingredient | CD | KD | BHB |
| --- | --- | --- | --- |
|  | TD.220115 | TD.220116.PWD | TD.230257 |
|  | g/kg | g/kg | g/kg |
| Casein | 100.0 | 180.0 | 100.0 |
| DL-Methionine | 1.6 | 2.88 | 1.6 |
| Corn Starch | 512.46 | 0 | 412.46 |
| Sucrose | 100.0 | 0 | 100.0 |
| Maltodextrin | 155.0 | 0 | 155.0 |
| Vegetable Shortening | 25.0 | 590.0 | 25.0 |
| Safflower Oil | 25.0 | 85.0 | 25.0 |
| 3-Hydroxybutyric Acid, liquid, customer supplied | 0 | 0 | 100.0 |
| Cellulose | 44.648 | 70.164 | 44.648 |
| Mineral Mix, w/o Ca & P (98057) | 13.4 | 24.8 | 13.4 |
| Calcium Phosphate, dibasic | 10.1 | 18.6 | 10.1 |
| Calcium Carbonate | 5.0 | 9.4 | 5.0 |
| Vitamin Mix, w/o choline, A, D, E (83171) | 5.0 | 13.875 | 5.0 |
| Choline Bitartrate | 2.5 | 4.625 | 2.5 |
| Vitamin E, DL-alpha tocopheryl acetate (500 IU/g) | 0.225 | 0.41 | 0.225 |
| Vitamin A, palmitate (500,000 IU/g) | 0.027 | 0.05 | 0.027 |
| Vitamin D3, cholecalciferol (50,000 IU/g in sucrose) | 0.03 | 0.056 | 0.03 |
| TBHQ, antioxidant | 0.01 | 0.14 | 0.01 |

TD.230257 is modified from TD.220115 to replace 10% cornstarch with 3-Hydroxybutyric Acid (BHB). The formula shows 10% inclusion, reflecting the net BHB addition. The preparation contains 50% water, so the actual inclusion of the ingredient is 20%. The 10% water from the BHB ingredient contributes to the water that is added in order to pellet the diet. BHB contains ~4.6 kcal/g. When 10% is added, BHB account for ~12% of total energy.
BHB = β-hydroxybutyrate supplemented diet, CD = control diet, KD = ketogenic diet.

**Table 2.** Dietary compositions. Values are expressed as a percentage of energy.

|  |  |  |  |
| --- | --- | --- | --- |
|  | Control diet | Ketogenic diet | BHB-supplemented diet |
| Energy (kcal/g) | 3.7 | 6.8 | 3.7 |
| Protein (E%) | 9.7 | 9.4 | 9.7 |
| Carbohydrate (E%) | 77.8 | 0.5 | 65.8 |
| Fat (E%) | 12.5 | 90.1 | 12.5 |
| BHB (E%) | 0 | 0 | 12 |

BHB = β-hydroxybutyrate.

DSS treatment was started on day 28 of the experiment by replacing regular drinking water with 2.5 % w/v DSS (40 kDa, TdB Labs, Uppsala, Sweden) solution (2.5 % w/v). From each diet group, eight animals were randomly allocated to DSS groups: DSS-treated control diet group (DSS-CD), ketogenic diet group (DSS-KD), and BHB-supplemented diet group (DSS-BHB) (Figure 1). DSS was administered for four days, after which regular water was given for two days before sacrifice. In addition to weight, food, and fluid consumption, consistency of stool, and presence of blood in the feces were monitored daily during this period. Based on these parameters, disease activity index (DAI) was determined by following a previously described scoring system (Wirtz et al., 2017) where scores from 0 to 4 are given for each of the following: weight loss, stool consistency and the degree of intestinal bleeding, and the final score for each animal is calculated as their average. If the animal reached the humane endpoint during the experiment, it was euthanized.

**Figure 1.**
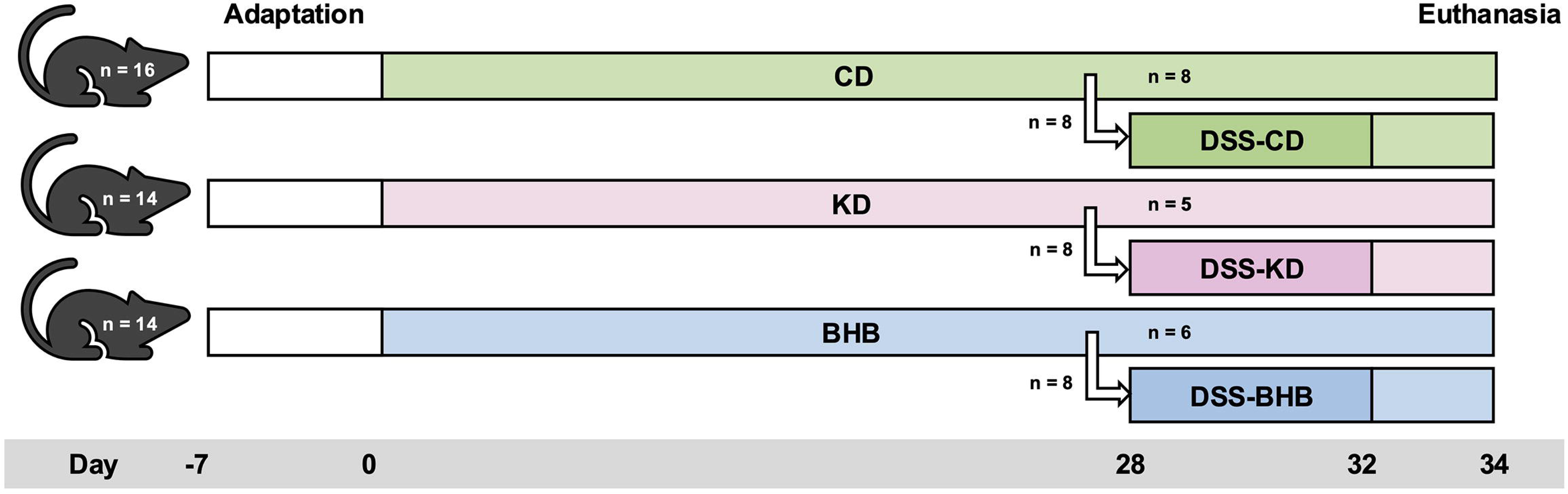
The set-up of the animal experiment.

### 2.2. Sample collection

Fecal samples were collected prior to DSS administration on day 27 of the experiment. For sample collection, animals were placed in empty cages and after 4 hours, fecal pellets were collected and frozen.

Animals were sacrificed under isoflurane (4 %, Vetflurane, Virbac, Carros, France) anesthesia by drawing blood from *vena cava* into EDTA-tubes (Kisker, Steinfurt, Germany). Blood samples were centrifuged at 2000 g for 15 min at 4 °C to separate plasma which was then frozen in liquid nitrogen.

The entire intestine was removed, and the length of the colon (from cecocolic orifice to rectum) measured. The colon was photographed for macroscopic evaluation before and after opening it longitudinally. Intestinal contents were collected, residues of which were flushed off with 0.9 % NaCl solution. Three 0.5-1 cm-long tissue sections were collected from the distal part of the colon. They were frozen in liquid nitrogen for biochemical analyses or fixed for histological analyses as described below. Samples were also collected from prematurely killed animals and used for analyses.

### 2.3. Intestinal permeability measurement

Fluorescein isothiocyanate–dextran (FITC–dextran) (4 kDa, TdB Labs) solution (600 mg/kg, 125 mg/ml) was administered to the animals 4 hours before the euthanasia via a gastric gavage. FITC- dextran concentration was analyzed from plasma diluted in PBS-T (136 mM NaCl, 8 mM Na2HPO4, 2.7 mM KCl, 4.46 mM KH2PO4, 0.1 % Tween, pH 7.4) with a fluorescence spectrophotometer at the excitation wavelength of 495 nm and the emission wavelength of 525 nm. The standard curve for determining the concentration in the samples was obtained by diluting known amounts of FITC– dextran in PBS-T.

### 2.4. Macroscopical evaluation

Photographs of colon were used to evaluate the extent of colitis based on three parameters: presence of diarrhea, visible fecal blood, and inflammation (edema and/or ulceration). Based on the severity, scores from 0-3 were given for each parameter by following a system described previously (Melgar et al., 2005). The scoring was performed blinded.

### 2.5. Histological analyses

Tissue sections from the most distal part of the colon were fixed in 4 % paraformaldehyde solution (Thermo Fisher Scientific, Waltham, MA, USA) for 36 h after which they were transferred to 70 % ethanol and stored at 4 °C. The fixed samples were cut into 2 halves to obtain 2 longitudinal pieces that were embedded in paraffin, sectioned at 4 μm thickness, and stained with hematoxylin and eosin (HE) dye.

The HE-stained slides were evaluated for histological changes and scored for severity of tissue damage and inflammation. The scoring followed a previously described system where separate scores from 0 to 3 for tissue damage and inflammation were summed to a combined score ranging from 0 to 6 (Wirtz et al., 2017). The scoring was modified due to the generally mild histological alterations observed in DSS-KD group. The mildest DSS-induced changes with minimal to mild mucosal damage received a tissue damage score of 0.5 and mild *lamina propria* mononuclear infiltrates a score of 0.5. The minimal to mild mucosal damage was defined as slight elongation and/or dilatation of the crypts and goblet cell hyperplasia as well as mild degenerative changes in the surface and crypt epithelium. Mucosal tissue damage (marked degeneration, flattening and/or dysplastic crypt epithelium) that did not include marked surface epithelial erosions or ulceration was scored as 1.5 and non-extensive damage extending beyond mucosa as 2.5. Inflammatory cell infiltration limited to the inner layer of *muscularis externa* received a score of 2.5. The scoring was performed blinded.

### 2.6. Biochemical analyses

Plasma BHB concentration was analyzed with a commercial enzymatic kit (Cayman Chemicals, Ann Arbor, MI, USA) after diluting the samples to the assay buffer. Levels of fecal calprotectin were analyzed with Mouse S100A8/S100A9 Heterodimer DuoSet ELISA (R&D Systems, Minneapolis, MN, US). Results were normalized against total protein determined with Pierce^TM^ BCA Protein Assay Kit (Thermo Fisher Scientific).

### 2.7. Targeted gene expression analysis

The expression of the TJ protein-coding genes *Cldn1, Cldn2, Cldn4,* and *Ocln;* and inflammatory marker genes *Tnf, Il1b, Il6,* and *Lcn2* in colonic tissue was analyzed with reverse transcription quantitative polymerase chain reaction (RT-qPCR) according to a protocol previously described (Toivio et al., 2023). In brief, RNA was extracted from colonic samples with NucleoSpin RNA Kit (Macherey Nagel, Duren, Germany). RNA concentration was determined, samples were diluted to the same concentration, and RNA was reverse transcribed to complementary DNA with iScript^TM^ cDNA Synthesis Kit (Bio-Rad). RT-qPCR was run with LightCycler® 480 SYBR Green Master (Roche Diagnostics Corp., Indianapolis, IN, USA) with the following amplification protocol: 10 min at 95 °C, 40 cycles of denaturation (15 s, 95 °C), annealing (30 s, 60 °C) and elongation (30 s, 72 °C). The primer sequences used are listed in Table 3. The results were calculated as relative quantities (RQ) of messenger RNA (mRNA) according to the Vandesompele method (Vandesompele J., 2002). The housekeeping genes used for normalization were *18S, Eef2,* and *Rplp0*.

**Table 3.**
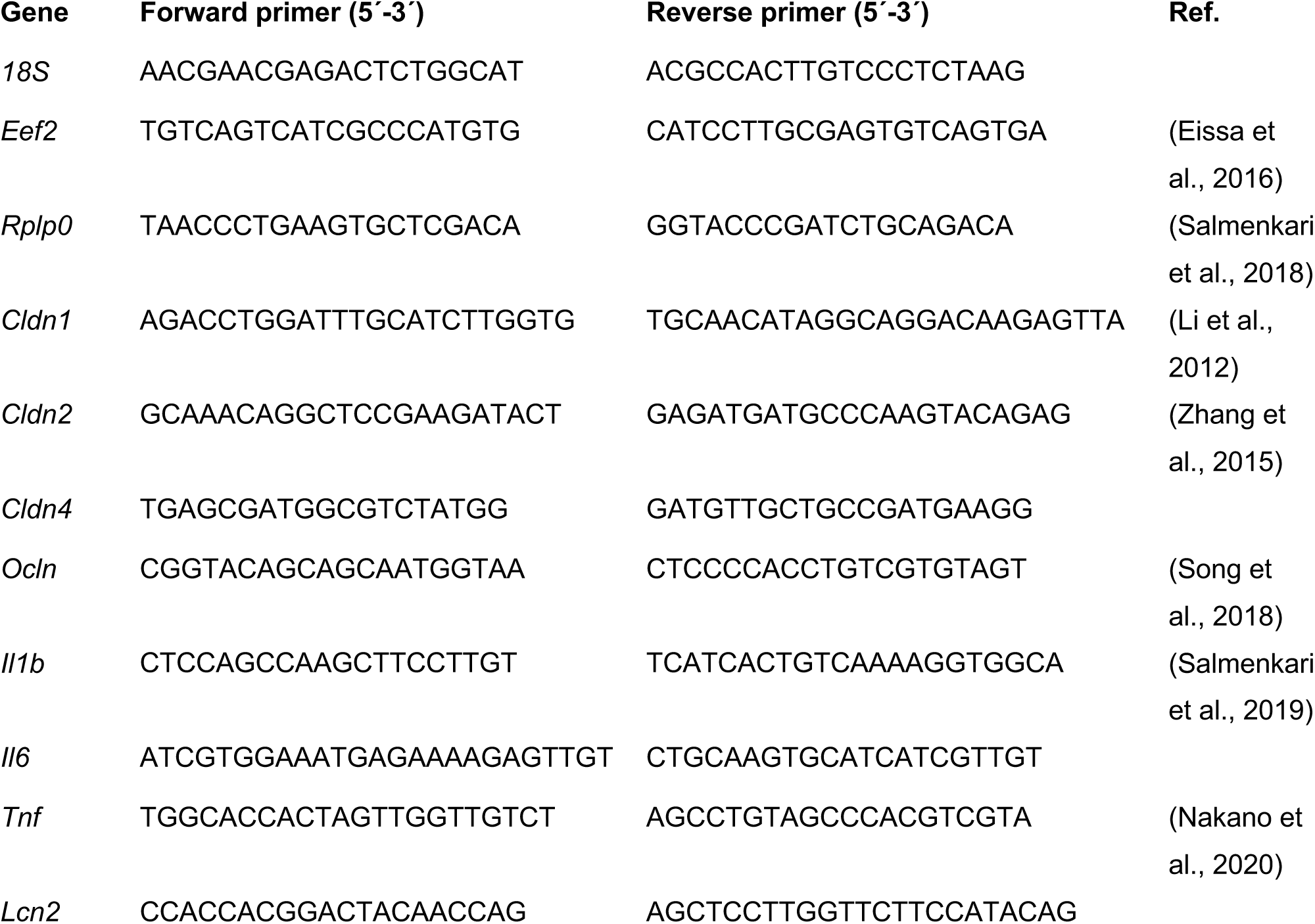
Primer sequences used in RT-qPCR analyses. If no reference is cited, the primer pair is designed by the authors.

### 2.8. Fecal microbiota analysis

The composition of the fecal microbiota prior to DSS administration was analyzed through 16S rRNA sequencing. A single fecal pellet was transferred to Lysing Matrix E tubes (MP Biomedicals, Santa Ana, California, USA) containing lysis buffer (MagMAX™, Thermo Fisher Scientific).

Samples were lysed using the TissueLyser II (Qiagen, Hilden, Germany), and the supernatant was transferred to a deep well block (Thermo Fisher Scientific). DNA extractions and purifications were carried out using the KingFisher Apex Purification System (Thermo Fisher Scientific) following the MagMAX™ Microbiome Ultra Nucleic Acid Isolation protocol. Paired-end DNA libraries with a 350 bp insert size were prepared using 16S rRNA V4 region primers (515F-806R) with dual indexing. Library quality was assessed with Tapestation, Qubit, and KapaQuant, and sequencing was performed using the MiSeq Reagent Kit v2 (500 cycles) on the Illumina MiSeq platform. The demultiplexed sequences were quality-controlled using DADA2. Microbiome bioinformatics, including alpha and beta diversity and relative abundance of specific taxa, were analyzed using QIIME2 (Bolyen et al., 2019), with differentially abundant taxa assessed using Analysis of Composition of Microbiomes with Bias Control (ANCOM-BC) (Lin & Peddada, 2020).

### 2.9. Statistical analyses

The data was analyzed, and figures created with GraphPad Prism 10 (Dotmatics, La Jolla, CA, USA). For fecal microbiota data, the figures were generated with QIIME2 (Bolyen et al., 2019). Mixed-effects model followed by Tukey’s multiple comparisons test was used for weight change and DAI. For other analyses, normal distribution was determined by Shapiro-Wilk test. Based normality, the data was analyzed either using one-way ANOVA followed by Tukey’s post-hoc test, or Kruskal-Wallis H test followed by Dunn’s post-hoc test. The level of statistical significance was set at p < 0.05. The data is illustrated as mean except for RT-qPCR results which are shown as geometric mean. In the analyses, each DSS group was compared to a healthy group on the same diet and all DSS groups were compared against each other. For microbiota data, pairwise Kruskal-Wallis H test was used for ɑ-diversity analyses (Shannon entropy score and Faith PD score) and pairwise Permanova test for β-diversity analyses (Unifrac analyses).

## 3. Results

### 3.1. Ketogenic diet alleviates DSS-induced macroscopic changes while BHB supplementation aggravates them

While all animals in DSS-CD and DSS-KD survived to the end of the experiment, in DSS-BHB two mice died and one had to be sacrificed prematurely. DSS administration resulted in significant weight loss which was ameliorated in DSS-KD group that did not lose weight (Figure 2A). DSS- BHB group exhibited a significant decrease in weight when compared to other DSS groups already on days two and three but the difference to DSS-CD lost significance on day four. DAI was lower in DSS-KD when compared to other DSS groups that exhibited visible signs of inflammation (Figure 2B). However, the difference was significant only between when DSS-KD and DSS-BHB. As expected, BHB levels were elevated in KD groups (Figure 2C). BHB supplementation resulted in only a small increase in plasma levels of the compound and the difference to animals on a control diet was not significant. DSS induced drastic macroscopic changes in the colon when compared to healthy controls in DSS-CD and DSS-BHB, whereas in DSS-KD there was no significant difference to KD (Figures 2D and 2E). While DSS-CD and DSS-BHB exhibited DSS-induced reduction in colon lengths, animals in DSS-KD were protected from colon shortening (Figures 2D and 2F).

**Figure 2.**
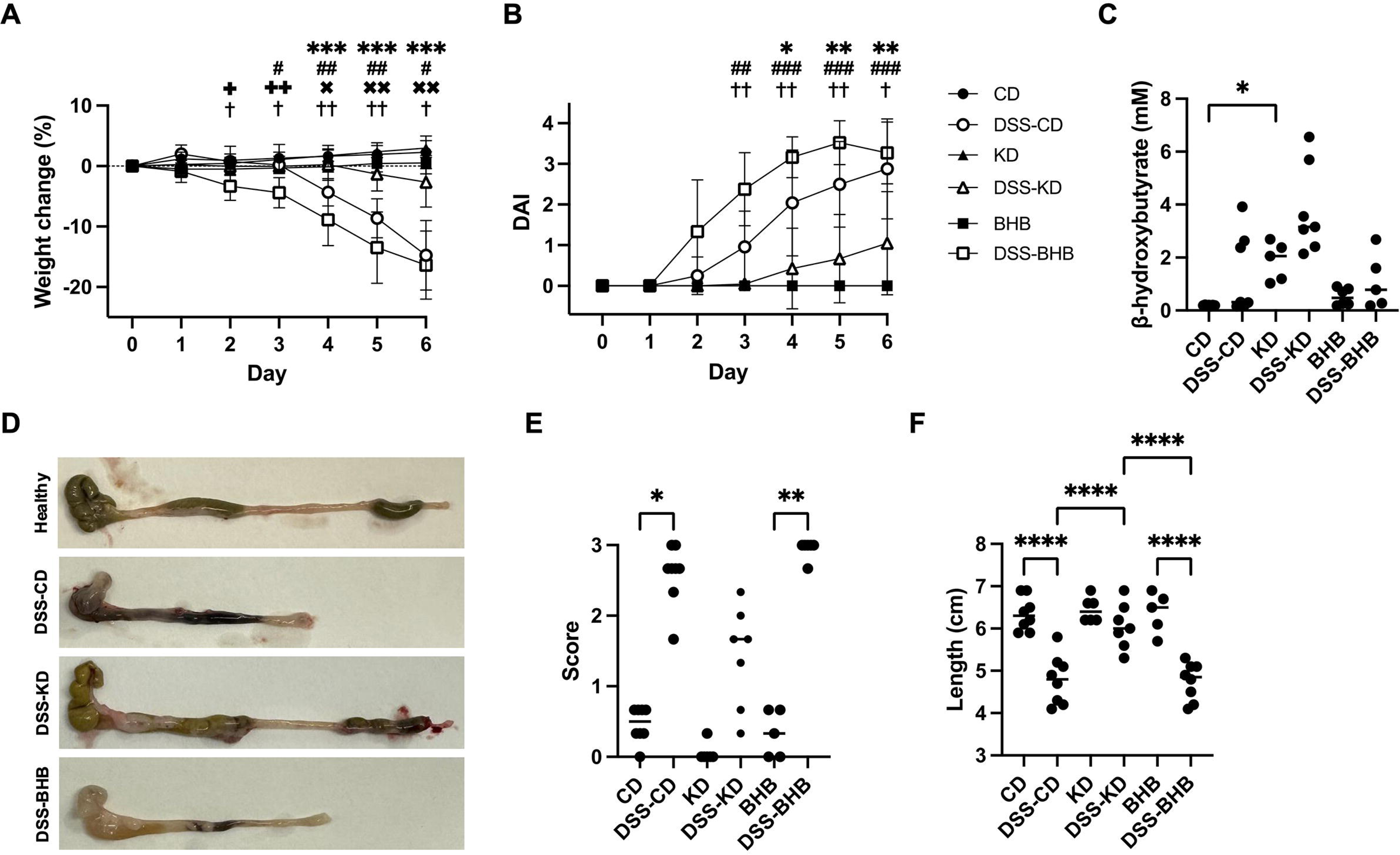
Macroscopic parameters of DSS-induced inflammation and plasma BHB levels. A. Weight change (mean ± SD, *n* = 6-8/group) as a percentage from the start of DSS administration demonstrating substantial weight loss in DSS-CD and DSS-BHB but not in DSS-KD. B. DAI scores (mean ± SD, *n* = 6-8/group) from the start of DSS administration showing substantial disease activity induced by DSS that was prevented by a ketogenic diet. C. Plasma β-hydroxybutyrate showing elevated levels upon a ketogenic diet but not β- hydroxybutyrate supplementation. D. Representative photographs of colons after DSS period showing marked macroscopic damage in DSS-CD and DSS-BHB but not in DSS-KD. E. Colitis score from the macroscopic evaluation of the colon suggestive of DSS-induced damage that is alleviated by a ketogenic diet. F. Length of the colon after DSS treatment showing significant shortening in DSS-CD and DSS-BHB but not in DSS-kD. Results for A. and B. are analyzed with mixed-effects model followed by Tukey’s multiple comparisons test, for C. and F. with Kruskal Wallis H test followed by Dunn’s post-hoc test, and for E. with one-way ANOVA followed by Tukey’s post-hoc test. In A. and B., * = CD vs. DSS-CD, # = BHB vs. DSS-BHB, x = DSS-CD vs. DSS-KD, + = DSS-CD vs. DSS-BHB, † = DSS-KD vs. DSS-BHB. * p < 0.05, ** p < 0.01, *** p < 0.001, **** < p 0.0001. DAI = disease activity index, DSS = dextran sodium sulfate, CD = healthy group with control diet, DSS-CD = DSS group with control diet, KD = healthy group with ketogenic diet, DSS-KD = DSS group with ketogenic diet, BHB = healthy group with β-hydroxybutyrate-supplemented diet, DSS-CD = DSS group with β- hydroxybutyrate-supplemented diet.

### 3.2. Ketogenic diet protects from DSS-related mucosal damage

DSS administration induced marked crypt loss, fibroplasia, and mucosal neovascularization, and in half of the samples, diffuse to focally extensively erosion of the surface epithelium with the remaining crypts having flat, degenerated, or dysplastic epithelium, and the surface epithelial cells showing marked degeneration and dysplasia (Figure 3). This was ameliorated in DSS-KD where the histopathological changes were minimal to moderate and mostly limited to the mucosa. Indeed, there was no statistically significant difference in tissue damage between KD and DSS-KD. On the other hand, in DSS-BHB, the histological alterations were qualitatively similar to those in DSS-CD but appeared to be more intense. Diffuse to focally extensive erosion of the surface epithelium was present in six of the seven samples. However, there was no significant difference in tissue damage score between DSS-CD and DSS-BHB.

**Figure 3.**
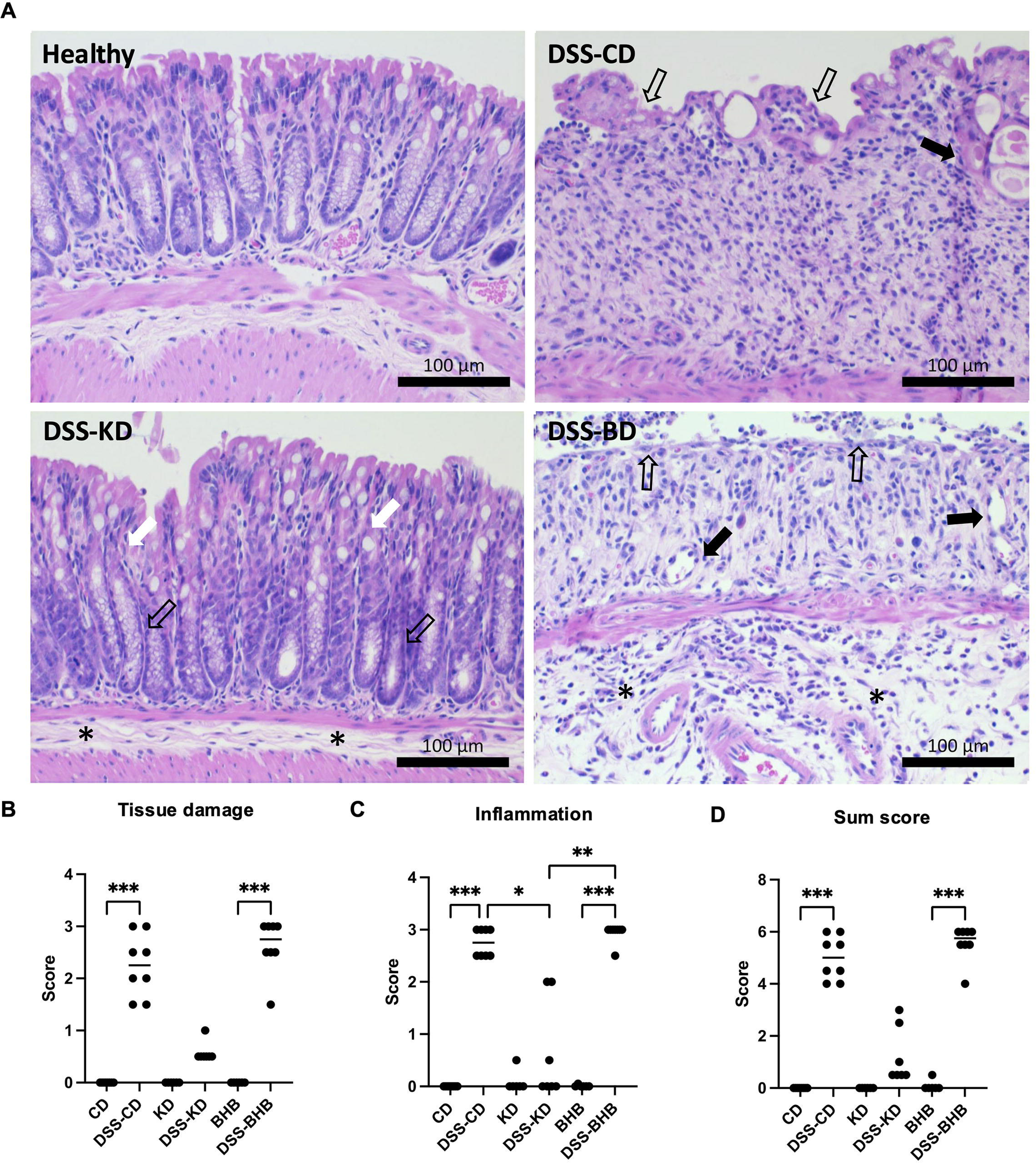
Histological findings of DSS-induced colitis. A. Microphotographs of hematoxylin-eosin-stained colon sections. Bars 100 µm. Control. No histopathological findings. DSS-CD. Crypt loss and lamina propria fibroplasia (astrisks). Surface epithelium (open arrows) and remaining crypt epithelium (arrow) show marked degeneration and dysplasia. DSS-KD. Minimal degenerative changes in the crypt epithelium and goblet cell hyperplasia (white arrows). Crypt elongation, epithelial lowering and basophilia point to crypt hyperplasia (open arrows). No cellular infiltrate in the submucosa (asterisks). DSS-BHB. The surface epithelium is eroded and covered with purulent infiltrate (open arrows). Lamina propria shows fibroplasia and apparent neovascularization (arrows) with plump endothelial cells. Moderate neutrophil and macrophage infiltration in the lamina propria and marked infiltration in the edematous submucosa (asterisks). B. Tissue damage score showing extensive damage in DSS-CD and DSS-BHB with little change in DSS-KD. C. Inflammation score demonstrating ample infiltration of inflammatory cells in DSS-CD and DSS-BHB which is ameliorated in DSS- KD. D. Combined sum score of tissue damage and inflammation indicating extensive DSS-induced histopathological changes in DSS-CD and DSS-BHB which are alleviated by ketogenic diet. Results are analyzed with Kruskal Wallis H test followed by Dunn’s post-hoc test. * p < 0.05, ** p < 0.01, *** p < 0.001. DSS = dextran sodium sulfate, CD = healthy group with control diet, DSS-CD = DSS group with control diet, KD = healthy group with ketogenic diet, DSS-KD = DSS group with ketogenic diet, BHB = healthy group with β-hydroxybutyrate-supplemented diet, DSS-CD = DSS group with β-hydroxybutyrate-supplemented diet.

In DSS-CD, moderate to marked mixed inflammatory cell infiltrate was present in the *lamina propria*. The submucosal inflammatory infiltrate consisted of macrophages and neutrophils. In most DSS-KD animals, there was no inflammatory infiltrate with three mice exhibiting minimal mononuclear cell or mixed cell inflammatory infiltrate. Again, the most severe changes appeared in DSS-BHB where several mice exhibited purulent infiltrate on the mucosal surface. Mixed inflammatory cell infiltrate filled the lamina propria and tissue damage extended throughout the intestinal wall in four mice. While no statistical difference was observed between DSS-CD and DSS-BHB, both groups exhibited a marked increase when compared to DSS-KD.

### 3.3. BHB supplementation worsens DSS-induced increase in inflammatory marker gene expression

As a marker of DSS-induced inflammation, we determined fecal calprotectin levels. All DSS groups displayed an increase in calprotectin levels when compared to their respective controls and differences were significant between KD and DSS-KD and BHB and DSS-BHB groups (Figure 4A). No differences between the DSS groups were found. We also analyzed the gene expression of inflammatory markers *Il1b*, *Il6*, *Tnf*, and *Lcn2* from colonic tissue. Based on the expression of these genes, DSS induced increased inflammation in all groups (Figure 4B). However, the elevation was more pronounced in DSS-CD and DSS-BHB which displayed statistically significant increases in *Il1b* and *Il6* when compared to CD and BHB groups, respectively. The level of inflammatory marker gene expression did not reach statistical difference in DSS-KD when compared to KD. Despite this, there were no significant differences in the expression these genes between DSS groups.

**Figure 4.**
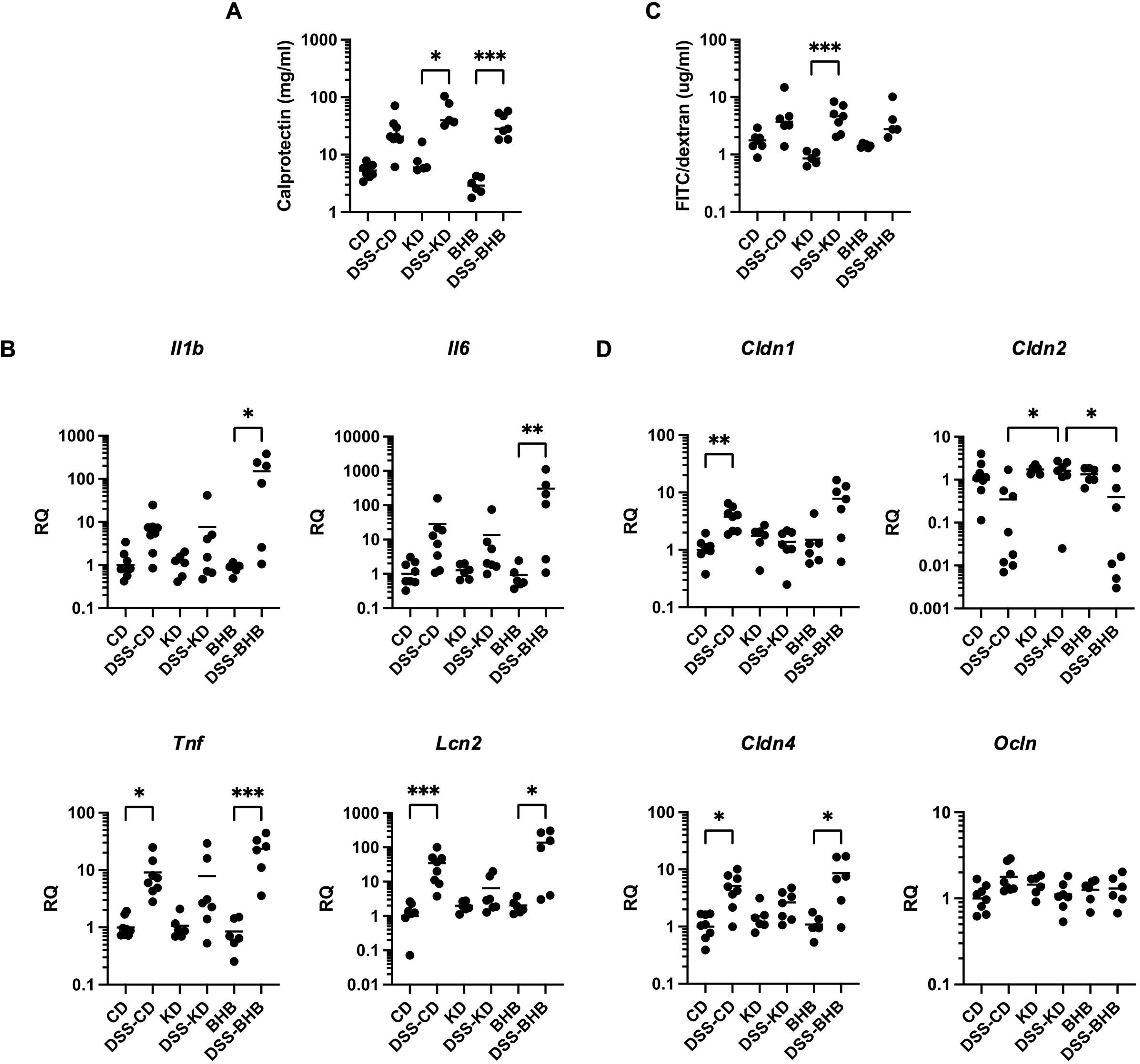
Selected markers of intestinal inflammation and permeability. A. Fecal calprotectin levels suggestive of a DSS-induced increase in all groups that is not prevented by a ketogenic diet. B Colonic mRNA expression of *Il1b*, *Il6*, *Tnf*, and *Lcn2* showing upregulation of transcription as a response to DSS to be aggravated in DSS- BHB group and alleviated in DSS-KD group. C. Intestinal permeability to FITC-dextran (4 kDa) measured as the plasma concentration of the compound indicating increased permeability upon DSS-induced inflammation. D. Colonic mRNA expression of *Cldn1*, *Cldn2*, *Cldn4*, and *Ocln* suggestive of DSS-induced changes in TJ protein expression that are mitigated with a ketogenic diet. Results are analyzed with Kruskal Wallis H test followed by Dunn’s post-hoc test. * p < 0.05, ** p < 0.01, *** p < 0.001. DSS = dextran sodium sulfate, CD = healthy group with control diet, DSS-CD = DSS group with control diet, KD = healthy group with ketogenic diet, DSS-KD = DSS group with ketogenic diet, BHB = healthy group with β-hydroxybutyrate-supplemented diet, DSS-CD = DSS group with β-hydroxybutyrate-supplemented diet.

### 3.4. Ketogenic diet normalizes the expression of tight junction protein-coding genes despite DSS-induced increased intestinal permeability

To assess intestinal permeability, the levels of orally administered FITC-dextran were analyzed from plasma. There was a trend towards lower permeability in healthy KD group when compared to other healthy groups. DSS induced an increase in FITC-dextran levels with all diets, indicating increased permeability, but the difference to their respective controls reached significance only in DSS-KD group (Figure 4C).

The expression of TJ-protein coding genes *Cldn1*, *Cldn2*, *Cldn4*, and *Ocln* was assessed to further evaluate barrier function (Figure 4D). In DSS-CD, the expression of *Cldn1* and *Cldn4* was elevated when compared to CD. *Cldn4* levels were also significantly higher in DSS-BHB when compared to BHB group. Both DSS-CD and DSS-BHB displayed a decrease in *Cldn2* when compared to DSS- KD. In DSS-KD, the expression of all TJ-protein coding genes was preserved at a level similar to healthy controls.

### 3.5. Ketogenic diet shifts the composition of fecal microbiota

The composition of the microbiota was analyzed from the three diet groups before DSS administration to investigate whether the diet-related changes in gut ecology might predict colitis severity. Shannon entropy score showed no differences in α-diversity between CD and BHB groups, but the score was higher for KD, indicating a higher overall microbial diversity based on richness and evenness (Figure 5A, Table S2). However, KD exhibited a lower Faith PD score compared to other groups, implying a less phylogenetically diverse microbiome (Figure 5B, Table S3). Unweighted Unifrac analysis showed significant differences in the composition of the microbial community between all groups, indicating different compositions even when the presence or absence of unique lineages is considered (Figures 5C, Table S4). Weighted Unifrac analysis revealed no significant difference between CD and BHB, suggesting that while there may be differences in the presence of specific lineages, these differences are not as pronounced when relative abundance of these lineages is considered (Figure 5D, Table S5). KD, however, differed from both groups, suggesting a distinct microbial community composition compared to both CD and BHB groups in terms of both lineage presence and their relative abundance.

**Figure 5.**
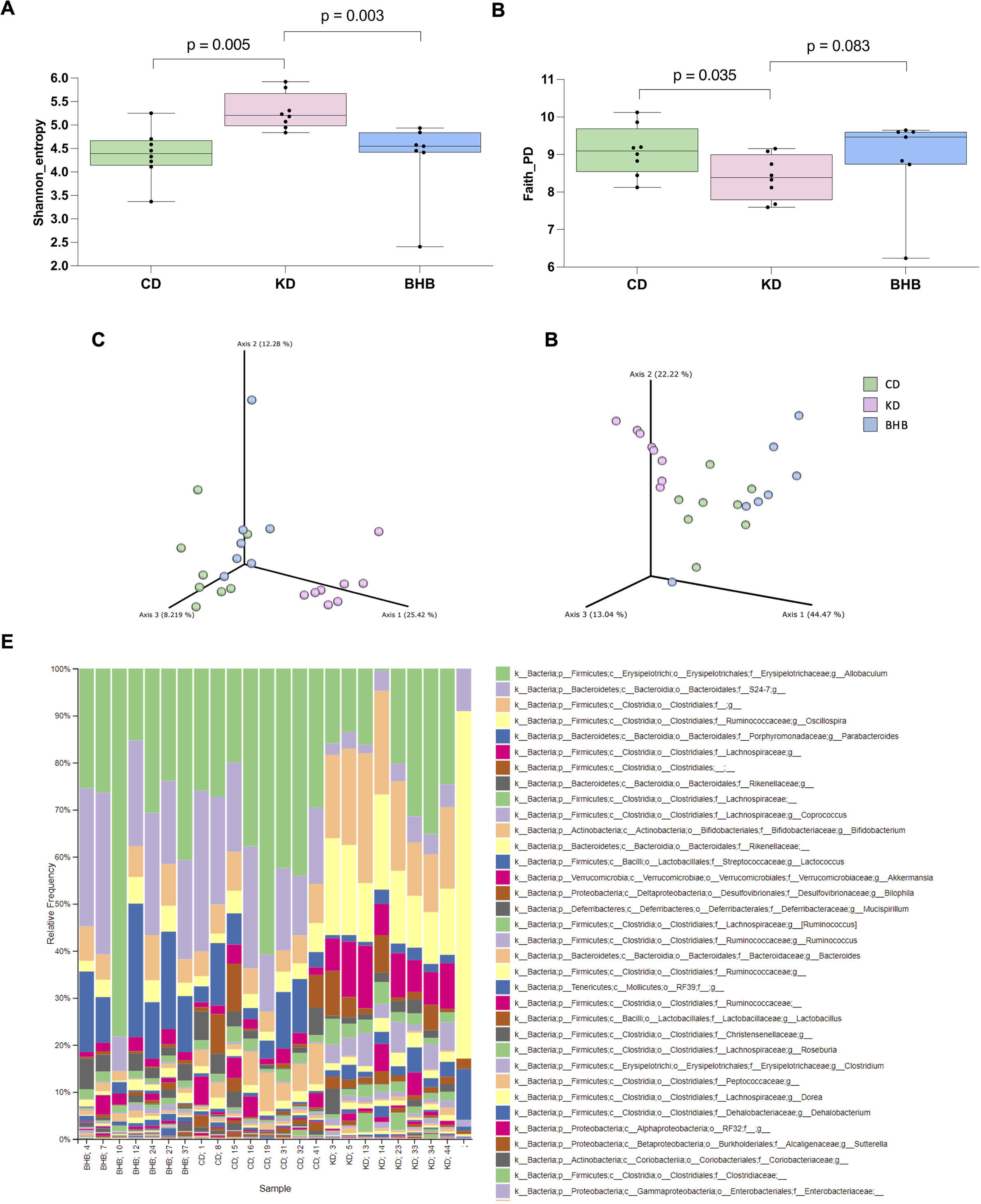
Diet-induced differences in fecal microbiota analyzed using 16S rRNA sequencing. A. Shannon diversity analysis indicating that KD group has a higher overall microbial diversity than CD and BHB groups. B. Faith phylogenetic diversity analysis suggesting KD group to have a gut microbiome that is less phylogenetically diverse than CD and BHB groups. C. Unweighted Unifrac dissimilarity analysis demonstrating differences in microbial community composition between the diet groups. D. Weighted Unifrac dissimilarity analysis showing that KD group has a distinct microbial community composition compared to CD and BHB groups. E. Bar plots of relative abundance of taxa in all samples showing notable differences in KD group compared to CD and BHB groups. Results for A. and B. are analyzed with Kruskal-Wallis H test and for C. and D. with Permanova test. CD = healthy group with control diet, KD = healthy group with ketogenic diet, BHB = healthy group with β-hydroxybutyrate-supplemented diet.

Figure 5E reveals notable differences in relative abundances of taxa between groups. While CD and BHB group resembled each other, there was a marked reduction in the relative abundance of taxonomic features associated with the genus *Allobaculum*, the family S24-7, and the genus *Parabacteroides* in KD group. Conversely, in KD group there was an increase in taxonomic features linked to the *Clostridiales* order, the *Oscillospira* genus, and the *Lachnospiraceae* family.

Discriminating features between groups were identified with ANCOM-BC analysis. Figure S1 shows the features that the relative abundance difference is larger than log102. In BHB group, *Enterococcus* was enriched whereas *Butyricoccus* and *Lactobacillus* were decreased when compared to both CD and KD (Figures S1A and S1C). *Anaeroplasma* and *Bifidobacterium* were also lower in comparison to CD and *Coprococcus*, *Lactococcus*, and *Dehalobacterium* less abundant when compared to KD. KD depleted *Sutterella* and *Bifidobacterium* and increased *Coprococcus* and *Defluviitalea*, when compared to both CD and BHB. Also, KD increased the abundance of *Clostridium*, *Dorea*, *Anaerotruncus*, *Bilophila*, and *Ruminococcus* when compared to CD (Figure S1B).

## 4. Discussion

We and others have previously found ketogenic diets to protect from DSS-induced colitis (Kong et al., 2021; Toivio et al., 2024). The aim of this study was to explore whether these protective effects could be achieved with exogenous ketone body supplementation alone as a more sustainable option to a ketogenic diet. Thus, we investigated the effect of high-dose supplementation with exogenous free acid BHB in the same model of colitis. We fed mice with three diets: a low-fat control diet with or without BHB or a ketogenic diet. After four weeks of feeding, we treated mice with DSS to induce colitis and assessed histological changes, intestinal permeability, tight junction protein levels, inflammatory markers, and microbiota composition. We found colitis and histopathological lesions to be alleviated by the ketogenic diet, but, against the hypothesis, the effects were not recapitulated by high-dose feeding of exogenous free acid BHB on a low-fat diet.

In mice fed the control diet, DSS administration resulted in weight loss, visible signs of colitis, colon shortening, and macroscopically detectable colonic damage. While mice in DSS-KD were protected from these changes, DSS-BHB mice were not. DSS-CD presented typical DSS-related histopathological lesions and damage alongside elevated inflammatory transcription of *Tnf* and *Lcn2* while the ketogenic diet alleviated these changes. This corroborates the earlier findings by us and others where ketogenic diets have protected from DSS-induced inflammation and damage (Abdelhady et al., 2023; Kong et al., 2021; Toivio et al., 2024). We saw DSS to induce a significant elevation in calprotectin levels in DSS-BHB and, interestingly, also in DSS-KD. Therefore, protective effects of the ketogenic diet may not extend to calprotectin. As opposed to the ketogenic diet, BHB administration did not protect the mice from colitis – in fact, BHB seemed to even aggravate inflammation and resulted in death or euthanasia of several mice in DSS-group earlier than planned. These findings are not in line with others who have reported BHB administration to alleviate DSS-induced histopathological damage in the colon when administered intraperitoneally (Abdelhady et al., 2023; Mohammed et al., 2024), rectally (Huang et al., 2022), or via a gastric gavage (Li et al., 2021) which suggest that the administration route, dose, and form of BHB, *i.e.* ester, salt, or free form, may determine the outcome of supplementation.

Despite a clearly milder colitis phenotype in KD mice, the DSS-induced intestinal permeability to FITC-dextran was similar in all diet groups. Interestingly, the permeability to FITC-dextran in healthy KD mice was lower than in CD or BHB groups. However, the ketogenic diet maintained the transcription of all measured TJ proteins on a level comparable to healthy animals. We saw DSS to induce an elevation in *Cldn4* levels in DSS-CD and DSS-BHB, whereas *Cldn2* expression was reduced in these groups. DSS-KD was protected from these changes. While claudin-4 is considered as a barrier-sealing TJ protein and claudin-2 pore-forming, we have previously observed similar changes in experimental colitis (Toivio et al., 2024). Claudin-4 protein expression is also elevated in IBD (Weber et al. 2008). Indeed, the increased expression of barrier-forming and the decreased expression of pore-forming TJ proteins may be a compensatory mechanism which could also explain why there were no differences in permeability to FITC-dextran between DSS groups.

Based on our study, high-dose continuous feeding with free acid in the context of a low-fat diet does not seem protect against DSS-induced colitis alike ketogenic diets, raising the question of the protective mechanism of KD. Ketone bodies themselves have anti-inflammatory properties (Youm et al., 2015). Here, as expected, the plasma levels of BHB were elevated in mice on the ketogenic diet, but in mice supplemented with BHB, the levels of the compound did not increase in plasma despite the significant dietary dose. In other colitis studies on exogenous BHB, plasma levels increased when the compound was administered intraperitoneally (Abdelhady et al., 2023; Mohammed et al., 2024) . In addition, feeding rats with a ketone monoester resulted in plasma BHB levels equal to those achieved by BHB injection and protected from DSS-induced colitis (Mohammed et al., 2024). A beneficial effect on DSS-colitis was also achieved with feeding mice with poly-D-3-hydroxybutyric acid, another BHB derivative which consumption increases plasma BHB levels (Suzuki et al., 2023). Thus, the levels of circulating ketones may influence the development of intestinal inflammation.

Development of intestinal inflammation and lesions is associated with gut microbiota, and therefore diet-induced changes in gut microbial composition can have significant consequences on the outcome of colitis. Indeed, microbiota seems to be a critical determinant of the severity of DSS- induced inflammation (Forster et al., 2022). In addition, the effects of KD on DSS-induced colitis seem to be related to changes in the composition of microbiota (Kong et al., 2021). In our study, the BHB group had a microbiota distinctly different from the KD group, which was protected from colitis, suggesting that changes in its composition may have mediated the differing outcomes. In the other mouse studies on BHB and experimental colitis, microbiota was not analyzed and thus, comparisons cannot be made. In rats administered DSS, KD and intraperitoneally administered BHB both mitigated decreases in *Fusobacterium* spp., and *Lactobacillus* spp. BHB also helped maintain *Clostridium* spp. levels (Abdelhady et al., 2023). It is possible that the different BHB form, dose, or administration regime may have modified the gut microbiota to a direction that worsened colitis in our setting. On the other hand, the protective effect of the ketogenic diet can also be mediated through the gut microbiota.

Our results contravene those of others who have reported BHB to alleviate DSS-induced colitis when administered via a gastric gavage (Li et al., 2021), or intraperitoneally (Abdelhady et al., 2023). In addition, BHB enemas seem to improve recovery from DSS-induced inflammation rectally (Huang et al., 2022). However, there are several differences between the studies which may explain the contradicting outcomes. Our mice had constant access to BHB-supplemented food whereas Li *et al*. (2021) orally administered BHB once a day. While the dose we used was determined based on previous literature, these studies were not done on models of intestinal inflammation: even though high-dose continuous feeding with diet containing 10-20 % exogenous ketones has shown benefit in rodent models of cancer (Poff et al., 2014) and epilepsy (Kovacs et al., 2019), a resembling dose seems to be detrimental in DSS-induced colitis based on our findings. In addition, we used free acid BHB which may have different effects from other forms of BHB. As opposed to BHB salts and esters, free acid R-BHB is molecularly identical to the endogenously produced BHB and in healthy humans, 10 g/d of the free form seems to be tolerated well (Pimentel-Suarez & Soto-Mota, 2023). However, our relative dose was orders of magnitude higher and administered to diseased animals. Overall, the contrast between our results and those of others highlights the importance of the administration route, dose, and form BHB in the context of intestinal inflammation. On the other hand, this study corroborates the previous findings on the benefits of a ketogenic diet in experimental colitis (Abdelhady et al., 2023; Kong et al., 2021; Toivio et al., 2024). This suggests that other KD-related metabolic changes or features of the diet, not only the increased supply of BHB, may be responsible for the protective effect of the diet.

## 5. Conclusions

While ketogenic diet ameliorates experimental DSS-induced colitis in mice, high-dose continuous feeding with free acid BHB seems to aggravate it. Ketogenic diet and BHB supplementation led to drastically different microbiota compositions which may partly explain the outcomes. Our results indicate that the protective effect of ketogenic diet might not be mediated by increased supply of BHB alone. Overall, replacing ketogenic diet with high-dose exogenous ketone supplementation might not offer benefit in the context of intestinal inflammation.

## Author contributions

Conceptualization, L.T., H.S., and R.K.; methodology, L.T., and H.S.; formal analysis, L.T., J.T., J.L., and K.L; investigation, L.T, J.T., J.L., and K.L. ; resources, R.K.; writing—original draft preparation, L.T and J.T.; writing—review and editing, J.L, K.L, M.L, H.S, and R.K.; visualization, L.T., J.T, J.L, and K.L; supervision, H.S, and R.K. All authors have read and agreed to the published version of the manuscript.

## Acknowledgment

We thank Heikki Vapaatalo for his help in designing the study and preparing the manuscript. We also thank the personnel of the Finnish Centre for Laboratory Animal Pathology, Helsinki Institute of Life Science, for the preparation and staining of the samples for histological analyses.

## Author declarations

None.

## Data availability statement

The data used in this study are available from the corresponding author upon reasonable request.

