## Supplementary materials for "Free acid β-hydroxybutyrate supplementation does not ameliorate dextran sodium sulfate-induced colitis similar to ketogenic diet in male mice"

**Table S1.** Pairwise Kruskal-Wallis test results for Shannon diversity analysis.

| **Group 1** | **Group 2** | **p-value** |
| --- | --- | --- |
| BHB (n = 7) | CD (n = 8) | 0.643 |
| BHB (n = 7) | KD (n = 8) | 0.003 |
| CD (n = 8) | KD (n = 8) | 0.005 |

**Table S2.** Pairwise Kruskal-Wallis test results for Faith Phylogenetic Diversity analysis.

| **Group 1** | **Group 2** | **p-value** |
| --- | --- | --- |
| BHB (n = 7) | CD (n = 8) | 0.908 |
| BHB (n = 7) | KD (n = 8) | 0.083 |
| CD (n = 8) | KD (n = 8) | 0.035 |

**Table S3.** Pairwise Permanova Test results for Unweighted Unifrac analysis.

| **Group 1** | **Group 2** | **Sample size** | **p-value** |
| --- | --- | --- | --- |
| BHB | CD | 15 | 0.006 |
| BHB | KD | 15 | 0.001 |
| CD | KD | 16 | 0.002 |

**Table S4.** Pairwise Permanova Test results for Weighted Unifrac analysis.

| **Group 1** | **Group 2** | **Sample size** | **p-value** |
| --- | --- | --- | --- |
| BHB | CD | 15 | 0.079 |
| BHB | KD | 15 | 0.001 |
| CD | KD | 16 | 0.002 |

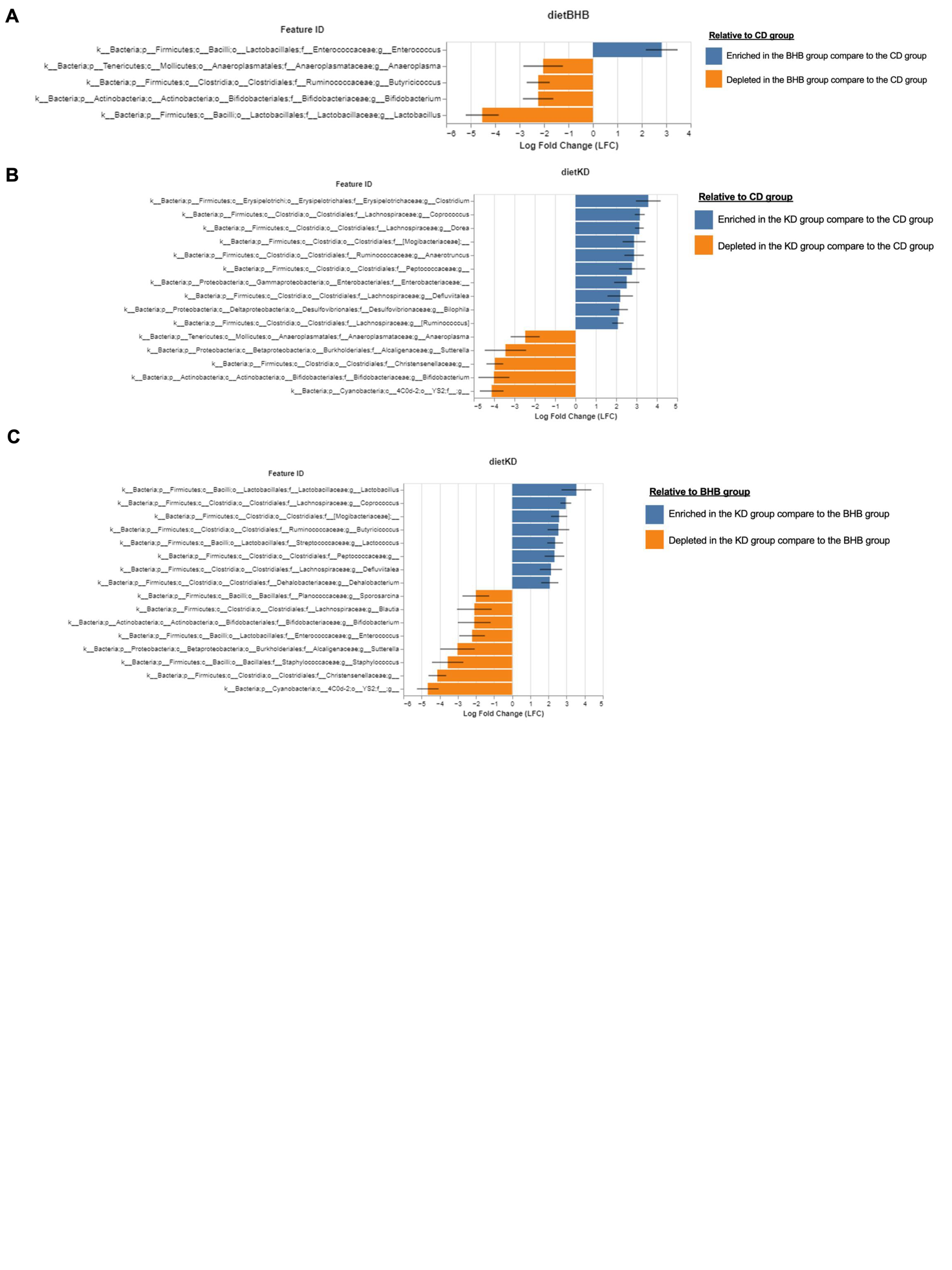

**Figure S1.** ANCOM-BC analysis data indicating significant differences in the relative abundance of specific microbial genera between groups. The graphs are showing the features that the relative abundance difference is larger than log_10_2 (p ≤ 0.05). A. CD vs. BHB, B. CD vs. KD, and C. KD vs. BHB. ANCOM-BC = Analysis of Composition of Microbiomes with Bias Control BHB = β-hydroxybutyrate supplemented diet group, CD = control diet group, KD = ketogenic diet group.
